# Effects of Large-Scale Oceanic Phenomena on Non-Cholera Vibriosis Incidence in the United States: Implications for Climate Change

**DOI:** 10.1101/528893

**Authors:** Chloë Logar-Henderson, Rebecca Ling, Ashleigh R. Tuite, David N. Fisman

## Abstract

**Purpose:** Epidemics of diarrhea caused by toxigenic strains of *Vibrio cholerae* are of global public health concern, but non-cholera *Vibrio* (NCV) species are also important causes of disease. These pathogens are thermophilic, and climate change could increase the risk of NCV infection. The El Niño Southern Oscillation (ENSO) is a “natural experiment” that may presage ocean warming effects on disease incidence.

**Method:** We obtained vibriosis case counts in the United States by digitizing annual reports from the U.S. Cholera and Other Vibrio Illness Surveillance system. Trends and environmental impacts (of ENSO and the North Atlantic Oscillation) were evaluated using negative binomial and distributed nonlinear lag models. Associations between latitude and changing risk were evaluated with meta-regression.

**Results:** Trend models demonstrated significant seasonality (P < 0.001) and a 7% annual increase in disease risk from 1999 to 2014 (annual IRR 1.071, 95% CI 1.061-1.081). Distributed lag models demonstrated increased vibriosis risk following ENSO conditions over the subsequent 12 months (integrated RR 1.940, 95% CI 1.298-2.901). The rate of change in vibriosis risk increased with state latitude (RR per 10° increase 1.066, 95% CI 1.027-1.107).

**Conclusion:** Vibriosis risk in the United States appears to be impacted by irregular large-scale ocean warming and exhibits a north-south gradient in rate of change as would be expected if changing disease incidence is attributable to ocean warming. Vulnerable populations, which include high-income countries with well-developed public health systems, may experience increased risk of this disease as a result of climate change.

## Introduction

Bacteria of the *Vibrio* genus are found in surface waters throughout the world and are responsible for causing several types of human infections, including cholera and non-cholera vibriosis [1]. While cholera is typically caused by toxigenic *Vibrio cholerae* serogroups O1 or O139, vibriosis is caused by non-toxigenic *V. cholerae* serogroups and roughly 12 other *Vibrio* species including *V. vulnificus, V. parahaemolyticus,* and *V. fluvialis* [2, 3]. While most individuals with vibriosis develop symptoms such as diarrhea and abdominal pain, some (particularly those with liver disease or immune compromise) may experience skin and soft tissue infection (typically characterized by bullous skin lesions) and septic shock [4]. Infection can be transmitted by food or by exposure to contaminated salt-water, and the most severely affected individuals often have immune compromise or liver disease [4]. Infections display summertime seasonality, which is believed to reflect the thermophilic nature of these bacteria, as well as increased leisure-related salt-water exposure and harvesting of seafood in the summer [1].

There has been an increase in non-cholera vibriosis reported in recent decades in the United States [2]. In 1988, the US Centers for Disease Control and Prevention (CDC) established the Cholera and Other *Vibrio* Illness Surveillance (COVIS) system, which initially focused on states (Texas, Alabama, Louisiana) forming part of the Gulf of Mexico coastline and expanded through the 1990s. By 2007, it was established as a national surveillance system at which time non-cholera vibriosis became nationally notifiable [2, 5]. The system relies on reports of laboratory-confirmed vibriosis cases from state and territorial public health authorities [2, 5].

The impact of large-scale climate phenomena, such as El Niño Southern Oscillation (ENSO), on cholera risk has been well-studied [6, 7]. The irregular and periodic nature of ENSO, and the similarity of ENSO-linked weather anomalies to those projected to occur with greater frequency with climate change, make ENSO a powerful natural experimental system for understanding the potential impact of climate change on infectious diseases and other health and economic issues [8]. Oceanic warming and extreme weather events including heat waves and hurricanes have also been associated with range expansion for vibriosis risk [9] and non-cholera vibrio outbreaks, respectively [10]. To our knowledge, the impact of ENSO and other large-scale climatic phenomena on vibriosis risk in the United States has not been previously examined.

We combined environmental exposure data with national and state-level vibriosis case counts derived from the COVIS system to evaluate the degree to which variation in vibriosis risk in the United States and in U.S. regions might be explained by large scale climatic phenomena such as ENSO. We also performed exploratory regional analyses to evaluate regional differences in evolving vibriosis risk, and to evaluate the possibility that changing patterns of risk are consistent with vibriosis range expansion.

## Methods

### Non-Cholera Vibriosis Data

Annual COVIS reports containing vibriosis case counts in the United States from 1997 to 2014 are publicly available, and were obtained from the United States Centre for Disease Control and Prevention [11]. The initial report, which included aggregate data for 1997 and 1998, was not used. Annual reported vibriosis case counts, by state, were extracted manually from published maps. Monthly national case counts were available from published graphs; we used the WebPlotDigitizer application (version 4.1) to accurately estimate case count numbers derived from these figures [12]. To ensure accuracy of extractions, monthly case count estimates were summed and compared to the reported annual national case counts. Incidence was estimated using U.S. national and state-level population denominators obtained from the U.S. Census Bureau [13]. Where necessary, estimates for inter-censal years were estimated via linear interpolation and extrapolation.

### Environmental Exposure Data

As noted above, ENSO is a complex climatic phenomenon manifested by changes in sea surface temperature and atmospheric pressure over the Pacific Ocean. ENSO has multiple attributes, including sea surface temperature, wind, air temperature, and cloud anomalies. For simplicity, we used the National Oceanic and Atmospheric Administration (NOAA) Multivariable ENSO Index (MEI) [14] as our measure of ENSO activity. The MEI is scaled from -3 (consistent with “La Niña”-like conditions) to +3, with high values corresponding to ENSO events. Monthly MEI values for January 1999 to December 2014 were obtained from the NOAA [14].

Although the effects of ENSO on North American weather are widespread, the strongest effects are observed in western states, and to some extent on Gulf states. Weather and ocean temperatures on the east coast of the United States are influenced by the North Atlantic Oscillation (NAO) [15], which manifests as fairly regular fluctuations in atmospheric pressure between Iceland and the Azores. The resultant pressure gradient influences land and sea surface temperatures, and precipitation patterns, in North America and Europe. Our measure of NAO strength, which we also obtained from NOAA, was the NAO Index. Like the MEI, this index is scaled from -3 to +3, with higher values associated with warmer temperatures and increased precipitation in the eastern United States and lower values associated with cooler, dryer conditions [15]. Unlike ENSO, which is defined in part by sea surface temperature anomalies, the relationship between sea surface temperature and NAO appears to be more complex. While certain analyses suggest an inverse relationship between NAO and sea surface temperatures at prolonged (4 year) lags [16], others suggest reversed causation, with NAO linked to preceding sea surface temperature anomalies in the Gulf Stream [17].

### Statistical Analysis

#### National Analyses

We evaluated year-on-year and seasonal trends in overallvibriosis incidence in the United States using count-based regression models that incorporated linear, quadratic, and cubic multi-annual trends, as well as Fast Fourier Transforms to account for seasonal oscillation. As both deviance and Pearson’s goodness-of-fit statistics suggested overdispersion of count data, negative binomial models were constructed. Census population estimates were used as model offsets.

Initial estimates of the lagged impact of ENSO and NAO on disease risk were generated by incorporating monthly MEI Index and NAO Index values into trend models at lags of 1 to 12 months. Models were constructed by backward elimination, with covariates retained for P < 0.2. Combined incidence rate ratios and standard errors for ENSO and NAO exposures at multiple lags were generated via linear combination [18].

The inclusion of multiple lagged exposures can result in several challenges, including difficulties with model interpretation, correlation in exposures at multiple lags, and non-parsimony of models. We therefore constructed distributed lag non-linear models to evaluate the integrated effects of environmental exposures at multiple lags [19]. These models characterize exposures as two-dimensional risk planes, referred to as “cross-bases”, and are defined as level of exposure by lag. We created two cross-bases, for MEI and NAO, at values ranging from -3 to +3, across 12 month lags, which were incorporated into generalized linear models that also adjusted for seasonality, and linear and cubic trends, using a log link function, in order to approximate negative binomial models constructed above. We assumed that ENSO and NAO effects on risk would be linear at any given lag, but modeled lag structure as a cubic polynomial. Both ENSO and NAO effects were evaluated at the uppermost extreme (+3) value, with index values of zero serving as referent.

#### Regional Analyses

COVIS reports annual vibriosis counts by state, with states grouped together according to the nature of their coastlines (Pacific, Gulf, Atlantic, or non-coastal). We evaluated relative incidence in each grouping using negative binomial models with state populations used as model offsets. We also evaluated linear trend terms from 1999 to 2014 for each region, and the associations between vibriosis risk and annual mean ENSO and NAO values. The non-coastal region was used as referent. Regional differences in trend terms, and the effect of average annual ENSO and NAO values, were evaluated using the Cochrane’s Q-statistic [20].

Linear trends in risk for each state were generated by estimating the annual incidence rate ratio for vibriosis by state. Negative binomial models for six states (Colorado, Iowa, Nebraska, New Mexico, South Carolina, and West Virginia) failed to converge. To evaluate whether trends might exhibit a north-south gradient suggestive of climate change effects, we constructed meta-regressive models using the latitude of state centroids as the explanatory variable [21]. Our hypothesis was that relative increases in risk due to climate change would be greater at northern than southern latitudes, particularly in coastal states, due to ocean warming.

All data used in this study are pre-collected aggregate counts and are publicly available. Negative binomial models and meta-regression models were constructed using Stata Intercooled version 15 (Stata Corp., College Station, Texas), while distributed lag non-linear models were constructed using the dlnm package for R, version 3.1.5 [22]. Datasets used for analyses can be obtained via Figshare at https://figshare.com/articles/Vibriosis_data_files/6856427.

## Results

From 1999 to 2014, a total of 10,800 cases of vibriosis were contained in COVIS reports, with an an additional 62 cases noted as part of a *V. parahaemolyticus* outbreak in Alaska in 2004. Of these 10,862 cases, we were able to assign 10,102 (93.0%) to a month of year in national-level data, while 10,857 (>99.9%) could be assigned to a specific state. Crude national incidence increased 3-fold during the period under observation, from 0.11 cases per 100,000 population in 1999 to 0.36 cases per 100,000 population in 2014, representing a 7% average annual increase in incidence (IRR 1.071, 95% CI 1.061 to 1.081). Significant seasonal oscillation was observed (P < 0.001) with a summertime predominance. As quadratic trend terms were not significantly associated with risk, and cubic trend terms were associated with a worsening of Akaike’s information criterion (AIC), the final trend model included only a linear term for year, as well as a Fast Fourier Transform (**Table 1**).

**Table 1.**
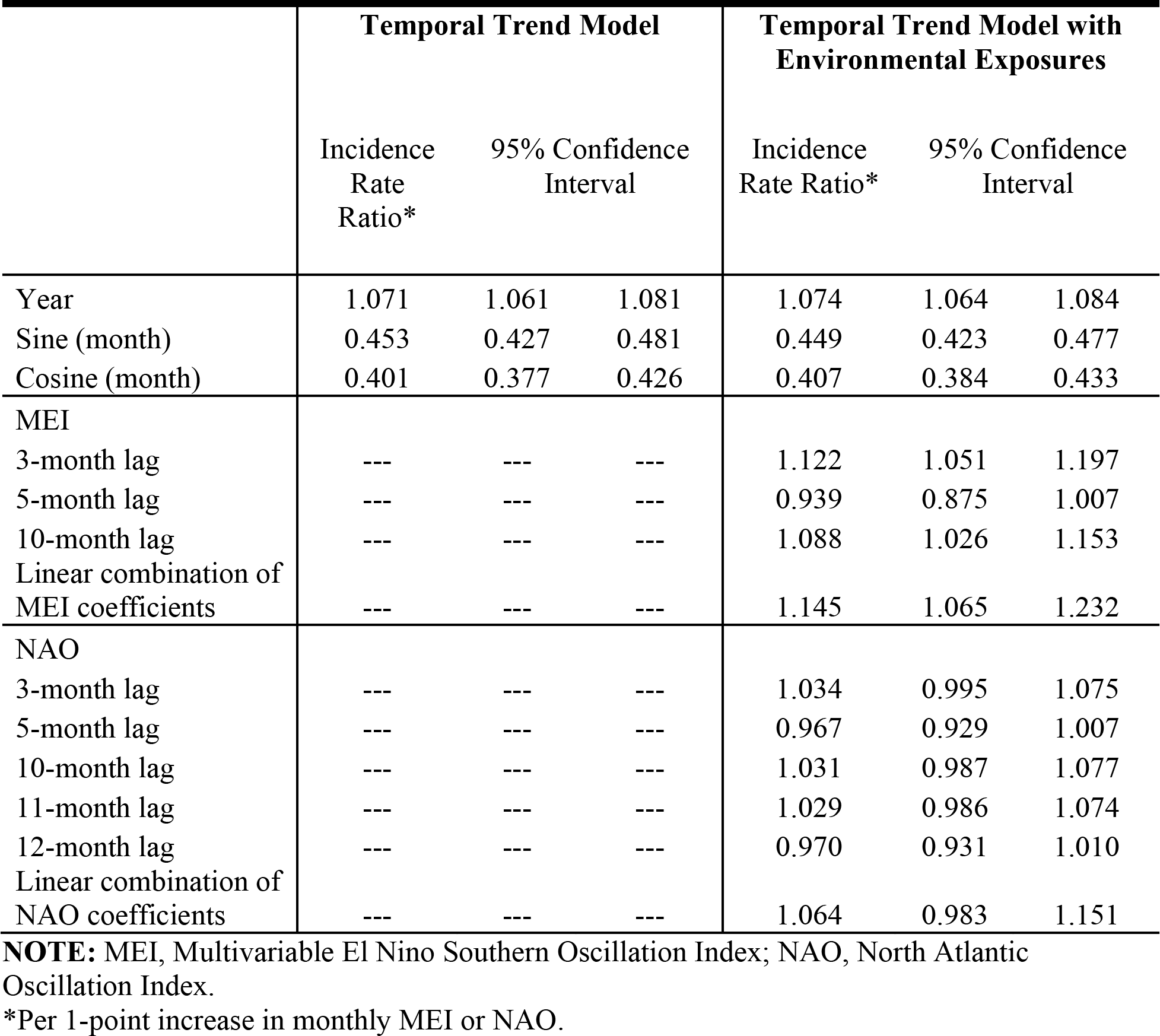
Temporal Trends and Impact of Environmental Exposures on National Vibriosis Incidence.

|  | Temporal Trend Model |  |  | Temporal Trend Model with Environmental Exposures |  |  |
| --- | --- | --- | --- | --- | --- | --- |
|  | Incidence Rate Ratio* | 95% Confidence Interval |  | Incidence Rate Ratio* | 95% Confidence Interval |  |
| Year | 1.071 | 1.061 | 1.081 | 1.074 | 1.064 | 1.084 |
| Sine (month) | 0.453 | 0.427 | 0.481 | 0.449 | 0.423 | 0.477 |
| Cosine (month) | 0.401 | 0.377 | 0.426 | 0.407 | 0.384 | 0.433 |
| MEI |  |  |  |  |  |  |
| 3-month lag | --- | --- | --- | 1.122 | 1.051 | 1.197 |
| 5-month lag | --- | --- | --- | 0.939 | 0.875 | 1.007 |
| 10-month lag | --- | --- | --- | 1.088 | 1.026 | 1.153 |
| Linear combination of MEI coefficients | --- | --- | --- | 1.145 | 1.065 | 1.232 |
| NAO |  |  |  |  |  |  |
| 3-month lag | --- | --- | --- | 1.034 | 0.995 | 1.075 |
| 5-month lag | --- | --- | --- | 0.967 | 0.929 | 1.007 |
| 10-month lag | --- | --- | --- | 1.031 | 0.987 | 1.077 |
| 11-month lag | --- | --- | --- | 1.029 | 0.986 | 1.074 |
| 12-month lag | --- | --- | --- | 0.970 | 0.931 | 1.010 |
| Linear combination of NAO coefficients | --- | --- | --- | 1.064 | 0.983 | 1.151 |

**NOTE:** MEI, Multivariable El Nino Southern Oscillation Index; NAO, North Atlantic Oscillation Index.
\*Per 1-point increase in monthly MEI or NAO.

ENSO was associated with enhanced risk of vibriosis at 3 and 10 month lags while ENSO at a 5 month lag was associated with attenuated risk. The overall effect of combined ENSO coefficients significantly increased vibriosis risk (combined IRR 1.145, 95% CI 1.064 to 1.231, P < 0.001). NAO effects were retained in the model at 5 different lags (3, 5, and 10-12 months), and were inconsistent in effect. Overall, the combined effect of NAO was non-significant (IRR 1.064, 95% CI 0.983 to 1.151, P = 0.12) (**Table 1, Figure 1**). Distributed lag non-linear models were also associated with a significant increase in vibriosis across lags of 1 to 12 months (integrated IRR 1.940, 95% CI 1.298 to 2.901). The integrated effect of NAO was again non-significant in distributed lag models (IRR 0.988, 95% CI 0.607 to 1.607) (**Figure 2**). As expected, regional analyses identified highest risks of infection in Pacific (IRR 13.412, 95% CI 10.607 to 16.959) and Gulf (IRR 7.752, 95% CI 6.167 to 9.744) regions, with elevated risk in the Atlantic region (IRR 4.125, 95% CI 3.491 to 4.873). Overall, yearly average ENSO was associated with a significant increase in vibriosis risk (IRR per 1 unit increase in mean MEI 1.169, 95% CI 1.012 to 1.350), while yearly average NAO was not associated with increased risk (**Table 2**).

**Table 2.** Temporal Trends, Regional Differences and Impact of Environmental Exposures on Annual Vibriosis Incidence in States.

|  | **Incidence Rate**<br>**Ratio\*** | **95% Confidence Interval** |  |
| --- | --- | --- | --- |
| Year | 1.108 | 1.090 | 1.126 |
| Yearly Average MEI | 1.169 | 1.012 | 1.350 |
| Yearly Average NAO | 0.909 | 0.758 | 1.091 |
| **Region** |  |  |  |
| Non-coastal | 1.000 | (referent) |  |
| Atlantic | 4.125 | 3.491 | 4.873 |
| Pacific | 13.412 | 10.607 | 16.960 |
| Gulf | 7.752 | 6.167 | 9.744 |

**NOTE:** MEI, Multivariable El Nino Southern Oscillation Index; NAO, North Atlantic Oscillation Index.
\*Per 1-point increase in yearly average MEI or NAO.

**Figure 1.**
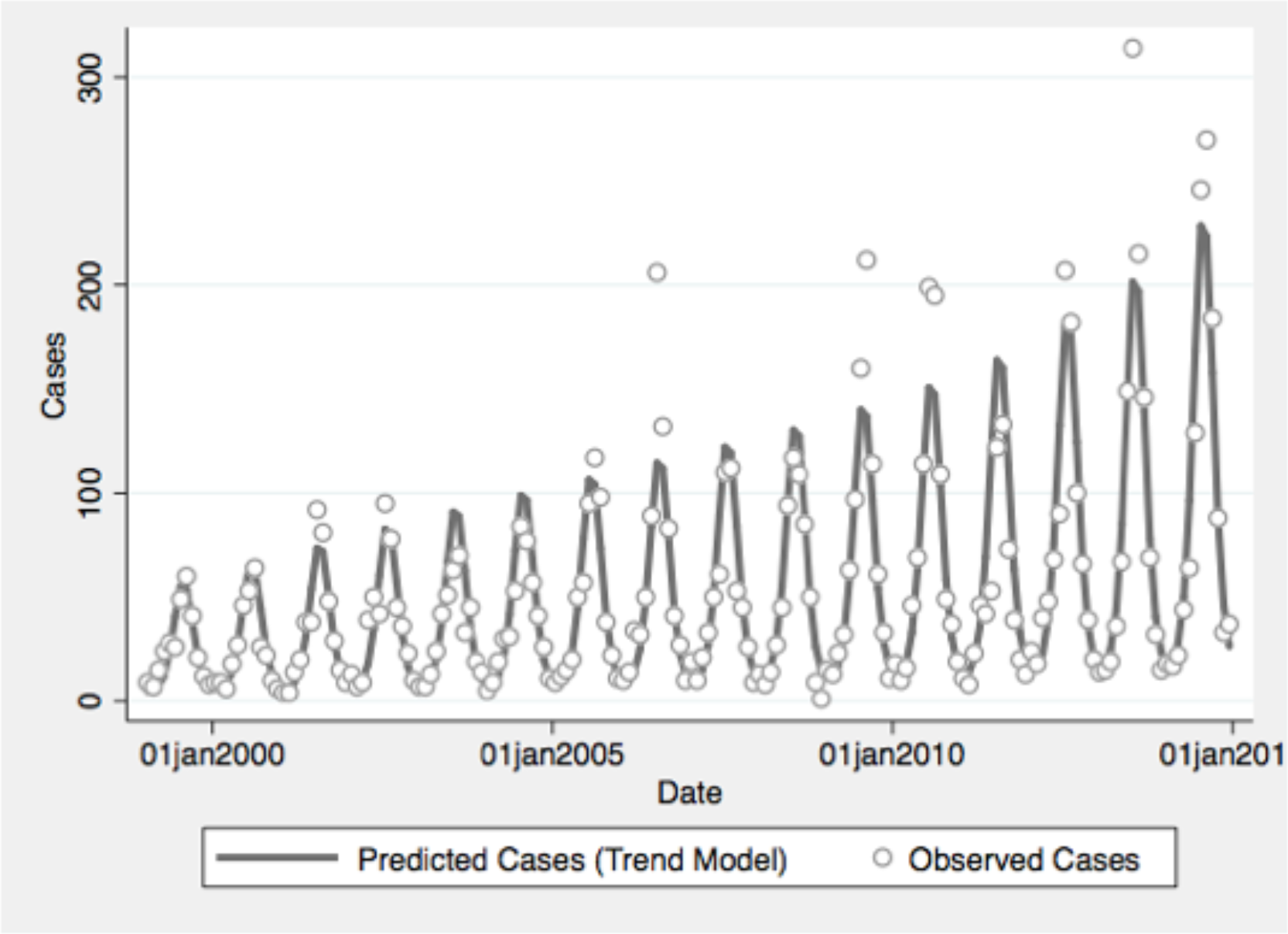
Reported and Model-Predicted Monthly Vibriosis Counts, United States 1999-2014. Observed counts are represented by circles; solid curve represents predictions from negative binomial model incorporating Fast Fourier Transform and year terms. Date is on the X-axis; counts are plotted on the Y-axis.

**Figure 2.**
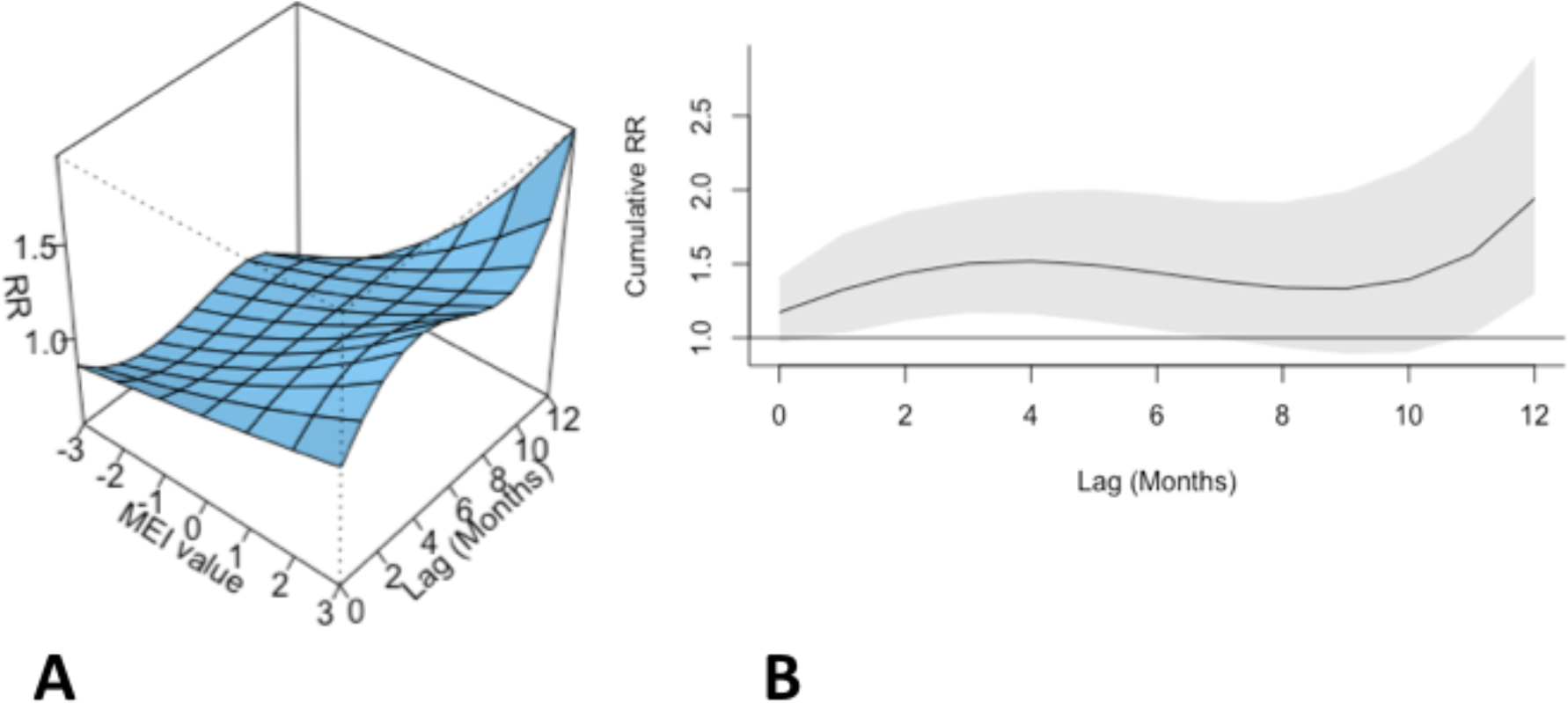
Association Between Lagged Multivariable El Niño Index and Vibriosis Risk, United States 1999-2014. (A) Risk surface represents modeled association between Multivariable El Niño Index (MEI) (scaled from -3, most La Niña-like, to +3, most El Niño-like, X-axis) and monthly vibriosis risk over 1-12 month lags (Y-axis). Associated relative risks are plotted on the Z-axis. (B) Cross-sectional relative risk associated with MEI of +3; lagged El Niño-like conditions are associated with downstream integrated relative risk of vibriosis (RR 1.940, 95% CI 1.298 to 2.900).

Significant increases in vibriosis risk were seen in non-Pacific regions during the period under observation, with significant heterogeneity in rate of change across regions (P for heterogeneity < 0.001); in the Pacific region, a non-significant increasing trend was observed (P = 0.067). The most marked increases in vibriosis risk were detected in the Atlantic and non-coastal regions (i.e., regions with the lowest incidence at baseline). In regional models, a 1 unit increase in mean MEI was associated with a 60% increase in vibriosis risk in the Pacific region, but no significant effect of ENSO on risk was was seen in other regions. No significant change in risk was seen with NAO in any region. Neither ENSO nor NAO effects were associated with significant regional heterogeneity (P for heterogeneity = 0.50 for ENSO; P = 0.52 for NAO) (**Table 3 and Figure 3**).

**Table 3.** Regional Models of Annual Vibriosis Incidence.

|  | Incidence<br>Rate Ratio* | 95% Confidence Interval |  |
| --- | --- | --- | --- |
| **Atlantic** |  |  |  |
| Year | 1.117 | 1.108 | 1.127 |
| Yearly Mean MEI | 1.032 | 0.959 | 1.111 |
| Yearly Mean NAO | 0.838 | 0.774 | 0.907 |
| **Pacific** |  |  |  |
| Year | 1.051 | 0.997 | 1.108 |
| Yearly Mean MEI | 1.601 | 1.031 | 2.486 |
| Yearly Mean NAO | 1.267 | 0.727 | 2.209 |
| **Gulf** |  |  |  |
| Year | 1.050 | 1.042 | 1.058 |
| Yearly Mean MEI | 1.016 | 0.948 | 1.089 |
| Yearly Mean NAO | 1.031 | 0.947 | 1.123 |
| **Non-coastal** |  |  |  |
| Year | 1.127 | 1.096 | 1.159 |
| Yearly Mean MEI | 1.174 | 0.916 | 1.505 |
| Yearly Mean NAO | 0.873 | 0.642 | 1.188 |

**NOTE:** MEI, Multivariable El Nino Southern Oscillation Index; NAO, North Atlantic Oscillation Index.
\*Per 1-point increase in yearly average MEI or NAO.

**Figure 3.**
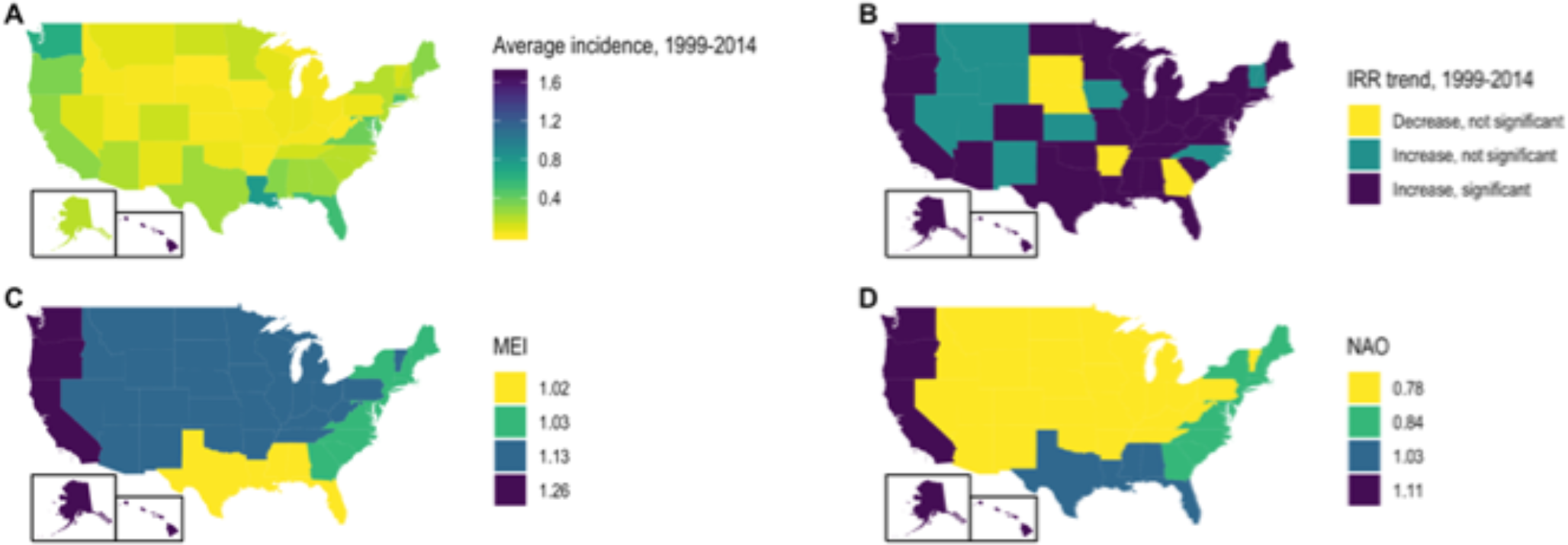
State-Level Incidence, Trends, and Environmental Influence on Annual Vibriosis Risk in U.S. States, 1999-2014. (A). Mean annual vibriosis incidence per 100,000 population, by state; (B) incidence rate ratios for year-on-year change in vibriosis incidence from negative binomial models, by state; (C) trends in vibriosis incidence from negative binomial models, by state; (D) incidence rate ratio for vibriosis risk with a 1-unit change in Multivariable El Niño Index (MEI), by region. Six grey-shaded states are those for which negative binomial models failed to converge.

State-level incidence rate ratios were also markedly heterogeneous (P < 0.001). In meta-regression models, increasing latitude was associated with the relative magnitude of increased risk (RR per 10° increase in latitude of state centroid 1.059, 95% CI 1.020 to 1.099). Adjustment for latitude resulted in a 36% reduction in between-study variance (τ^2^) [23]. There was no significant association between region and rate of change, after adjustment for latitude. We also found no significant interaction between latitude and whether or not a state was coastal (P for multiplicative interaction term = 0.87) (**Figure 4**).

**Figure 4.**
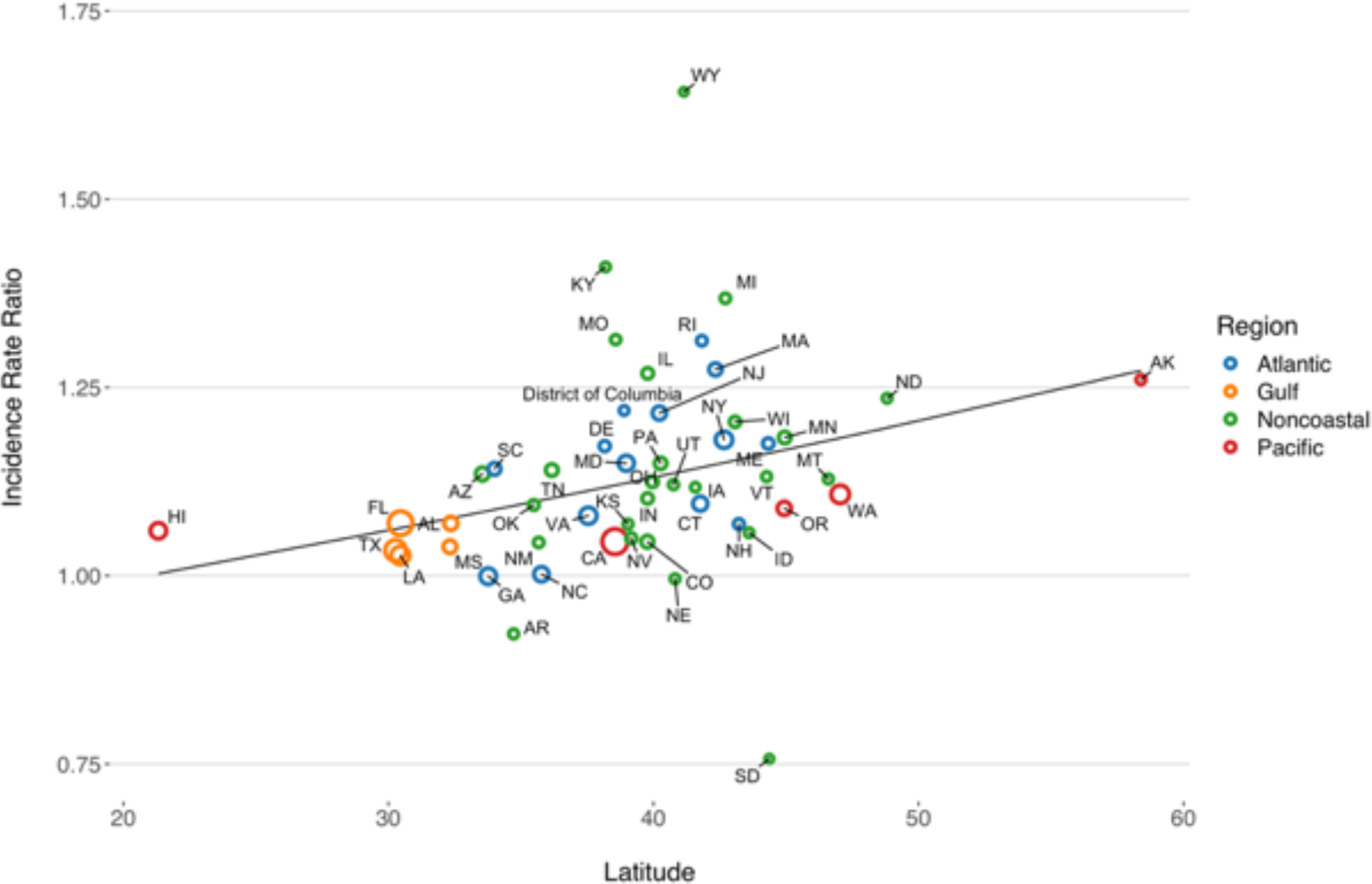
Association Between Latitude and Linear Trend in Vibriosis Incidence, United States 1999-2014. Correlation between state latitude and average yearly incidence rate ratio (IRR) for vibriosis. Each circle represents a single US state or the District of Columbia, with size inversely proportional to variance in IRR estimates, corresponding to the weight assigned to each state. Circles are colour coded according to COVIS regions. Fitted lines represents the association between state latitude and IRR as predicted using univariable meta-regression (relative change in IRR per 10° increase in latitude of state centroid 1.059, 95% CI 1.020 to 1.099). Note that six states for which negative binomial models did not converge are excluded.

## Discussion

Given that toxigenic *V. cholerae* is a sentinel global pathogen due to its virulence, capacity for genetic reassortment, and epidemic potential, the ecology and dynamics of this pathogen have been extensively studied [7, 24, 25]. However, other *Vibrio* species of public health importance can also cause severe human disease, and their associated disease burden appears to be increasing. An increase in disease incidence may reflect increased pathogen abundance in salt water environments, as well as an increase in the prevalence of immune compromised states in the human population, as individuals with immune compromise or iron overload states appear to be at greatest risk for severe vibriosis [2]. Due to the thermophilic nature of *Vibrio* species and their increased abundance in warmer waters with decreased salinity, there is reason to anticipate that *Vibrio* infection incidence will increase in coming decades as a result of climate change-associated ocean warming [9].

Using publicly available *Vibrios* case count data from the United States COVIS system, we evaluated vibriosis trends and their relation to large scale oceanic phenomena. We also examined the association bewtween rate of increase and latitude, as we hypothesized that relative increases in risk would be greatest at northernmost latitudes where vibriosis has historically been less common. Our observations suggest that climate change-driven range expansion may be in progress for this disease, and the sensitivity of vibriosis incidence to El Niño-like conditions suggests that climate change is likely to drive further increases in vibriosis incidence in the future. After adjusting for long-term trends and seasonal oscillation, we found vibriosis incidence in the United States to be very sensitive to the ENSO. Using distributed lag non-linear models to examine risk effects at multiple lags, we found that vibriosis incidence in the United States is expected to double in the year after strong ENSO-like conditions (Multivariable ENSO Index = 3). This finding is consistent with existing work on the ecology of cholera in low- and middle-income countries [6, 26]. The regional analyses suggested these effects are concentrated in the Pacific region, where states are most strongly teleconnected to ENSO and where baseline vibriosis incidence is highest.

Strong ENSO effects may provide insight into future changes in disease epidemiology that may occur under climate change scenarios such as increases in temperature, precipitation anomalies, and ocean warming effects [8]. As current anthropogenic climate change is a global phenomenon with no recent precedent, it has been noted that the irregular nature of ENSO allows it to serve as a natural experiment for exploring future climate change effects [8]. The mechanisms whereby ENSO might increase vibriosis risk are likely to be complex; however, conditions like changes in salinity, ocean warming, and increased abundance of nutrients and flow of freshwater into oceans are all associated with ENSO [27, 28], as well as with *Vibrio* abundance [29, 30]. It should be noted that, as with *V. cholerae,* non-cholera *Vibrios* have complex ecology, and their abundance may be impacted by the abundance of the cyanobacteria and dinoflagelates with which they are associated; these microbial populations are also sensitive to changes in nutrients, water temperature and salinity such as those that accompany ENSO [28, 31]. The complexity of *Vibrio* ecology, shown by differing effects of ENSO at multiple lags due to its impact on different components of the ecosystem, make the distributed lag approach we have applied particularly attractive.

Since weather and sea surface warming in the Pacific region are more likely to be influenced by ENSO than the Atlantic region of the United States, we also explored the impact of the NAO on vibriosis incidence. We hypothesized that NAO would be associated with regional vibriosis incidence in the Atlantic and possibly the Gulf regions. However, compared to the large effects observed with ENSO at the national leveland in the Pacific region, the effects associated with NAO were inconsistent in national analyses, and absent in regional analyses. Given that NAO is predominantly a pressure phenomenon (rather than a phenomenon explicitly related to ocean temperature like ENSO), the absence of clear and consistent links to vibriosis may be unsurprising.

Range expansion of vibriosis has been reported in Northern Europe and appears to correlate with increasing Baltic Sea temperatures [9]. We therefore sought to evaluate the possibility that there was a north-south gradient in rate of change in vibriosis risk in the United States. The meta-regression approach we applied here was identical to the approach applied in our earlier work on Lyme disease [21]. The gradient we observed would be consistent with range expansion for vibriosis risk as a result of warmer oceans at higher latitudes. However, other mechanisms (such as increased reporting in jurisdictions not previously believed to be risk-areas for vibriosis) could produce similar patterns. This is possible since the COVIS system has seen increased coverage and increased case submission over time, particularly once non-cholera vibriosis became nationally notifiable in 2007 [5]. However, given that similar trends in rates of vibriosis have been reported using FoodNet data, which involve active rather than passive surveillance, it is unlikely that these trends reflect reporting artifact [2]. Furthermore, an ocean warming effect would be consistent with the ENSO effects seen nationally, and in the Pacific region, in our earlier analyses.

Our analysis is inevitably subject to limitations. As we only have access to case counts by time period and region, we cannot account for changing reporting behavior as a driver for observed trends. However, it should be noted that we do account for such temporal trends in our models, and the ENSO effects we observe are robust after accounting for trends, regardless of their underlying mechanism. Our lack of access to individual case data means that we cannot perform nuanced subgroup analyses that consider case comorbidity or exposure history.

Extracting case counts from graphical plots produced counts similar to those contained in reports, but is likely to have resulted in some random misclassification of these counts; while this may have resulted in diminished statistical power, such misclassification would not have resulted in spurious associations or systematic bias in effects.

In summary, we used publicly available data on vibriosis incidence in the United States to perform the first (to our knowledge) evaluation of the possible contributions of ongoing environmental change to observed increases in vibriosis incidence in North America. Broadly, we found two lines of evidence suggesting that climatic change and ocean warming may be important drivers of observed increases in risk. First, *Vibrio* risk was strongly associated with lagged ENSO-like oceanic changes, and second, a north-south gradient in relative increase in risk. Attributing such changes to ocean warming and climatic change is biologically plausible, and consistent with effects observed in other geographic locales with *V. cholerae*. Our work suggests that vibriosis may be an important sentinel for infectious disease impacts of climate change, and our methods also present a model that can be applied to other infectious diseases. Anticipating future trends in vibriosis as oceans warm may help with projections of likely future burden of disease, and may help guide and prioritize preventive strategies.

